## Supplemental Appendix for "Active surface flows accelerate the coarsening of lipid membrane domains"

### I. MOVIE DESCRIPTION

**Movie S1:** Passive domain coarsening is observed experimentally on a phase-separated lipid bilayer. A single-phase planar lipid membrane was previously heated to ≈ 37°C, and is shown here as it cools to room temperature. Liquid-ordered domains (green) nucleate and coarsen as the membrane cools in the absence of flow.

**Movie S2:** Actomyosin-driven flows accelerate lipid domain coarsening on an experimental planar bilayer. (*Upper left*) Liquid-ordered domains (green) are initially arrested by actin, but rapidly change morphology and grow when activity is triggered at $t = 0$. (*Upper right*) An actomyosin cortex adsorbed to the lipid membrane via electrostatic attractions begins to contract when adenosine triphosphate (ATP) is introduced at $t = 0$. Actin is pulled inward toward a central cluster as it contracts. This actin cluster appears to delaminate from the surface allowing domains to coalesce in its wake. (*Lower*) Actin (magenta) and lipid domain (green) images are superimposed upon one another.

**Movie S3:** Numerical simulation of Cahn-Hilliard model for passive coarsening. The model parameters are ($M$, $\kappa$)=(1, 0.25).

**Movie S4:** Numerical simulations of Cahn-Hilliard model with experimentally obtained actin flow fields at Pe $= 10^{-2}$ (*left*), Pe $= 5 \times 10^{-3}$ (*center*), and Pe $= 10^{-4}$ (*right*). The model parameters are ($M$, $\kappa$)=(300, 16).

**Movie S5:** Numerical simulations of Cahn-Hilliard model with shear flow (*left*), and rotational flow (*right*). The model parameters are ($M$, $\kappa$, $\dot{\gamma}$)=(1, 0.25, 0.04), which correspond to Pe $= 10^{-2}$.

In movies S3-5, the simulation timescale is non-dimensionalized by the characteristic time scale of the Cahn-Hilliard model, $t_c = \kappa/M$. The timestamps in movies S3 and S4 are scaled by 10 to match with the timescale of the experiments.

### II. DETAILED EXPERIMENTAL METHODS

#### A. Buffers

Filamentous actin buffer (F-buffer) consists of 50 mM Tris (pH 7.5), 2 mM magnesium chloride, 0.5 mM adenosine triphosphate (ATP), 0.2 mM calcium chloride, 25 mM potassium chloride, and 1 mM dithiothreitol (DTT). DTT was added to all buffers immediately before use to preserve its reactivity.

Assay buffer (A-buffer) consists of 25 mM imidazole (pH 7.4), 4 mM magnesium chloride, 1 mM (ethylene glycol-bis(β-aminoethyl ether)-N,N,N',N'-tetraacetic acid) (EGTA), 25 mM potassium chloride, and 1 mM DTT.

Globular actin buffer (G-buffer) consists of 2 mM Tris (pH 8.0), 0.2 mM calcium chloride, 0.5 mM DTT, 1 mM sodium azide, and 0.2 mM ATP.

#### B. Actin and myosin preparation

Rabbit skeletal muscle actin was purified from muscle acetone powder (Pel-Freez, catalog no: 41995-2, Lot 16743) using standard methods [1, 2]. No rabbits or other animals were directly involved in this study. Actin was stored as depolymerized globular actin (G-actin) at -80°C in G-buffer with 6% sucrose until use.

---

* These authors contributed equally to this work

†

Actin was labeled with fluorescent Alexa Fluor 555 NHS Ester (Succinimidyl Ester) (Invitrogen catalog no: A20009) for microscopic visualization. G-actin was reacted with NHS-Alexa Fluor 555 in HEPES buffer at room temperature for 30 minutes. 2x-concentrated F-buffer was then added to the G-actin, quenching the NHS reaction and causing G-actin to polymerize to F-actin. F-actin polymerization proceeded for 30 minutes at room temperature, and then overnight at 4°C. Labeled F-actin was centrifuged at 142,000 × g for 30 minutes, and the pellet collected. Unreacted dye and defective G-actin monomers and oligomers that were unable to polymerize were discarded in the supernatant. Labeled F-actin was dissolved in G-buffer, and allowed to de-polymerize for three days at 4°C before freezing and storing in 6% sucrose at -80°C.

Rabbit skeletal muscle myosin II and heavy meromyosin (HMM) were purified from rabbit skeletal muscle (Pel-Freez Biologicals, Rogers, Arkansas) using standard methods [3]. Myosin II was frozen in 150 mM potassium phosphate buffer (pH 7.5) with 6% sucrose and 10 mM EDTA at -80°C until use. HMM was frozen in 10 mM potassium phosphate buffer (pH 7.0), 100 mM potassium chloride, 0.3 mM EGTA, 1 mM DTT, and 6% sucrose at -80°C until use.

#### C. Giant unilamellar vesicle (GUV) preparation

Giant unilamellar vesicles (GUVs) were prepared using the established method electrofomation [4]. Briefly, lipids were mixed with the following composition: 44.7% 1,2-dioleoyl-sn-glycero-3-phosphocholine (DOPC, Avanti catalog no: 850375P), 34.7% 1,2-dipalmitoyl-sn-glycero-3-phosphocholine (Avanti catalog no: 850355C), 15% cholesterol (TCI Chemical, catalog no: C3624), 5% 1,2-dioleoyl-3-trimethylammonium-propane (DOTAP, Avanti catalog no: 890890P), 0.3% 1,2-distearoyl-sn-glycero-3-phosphoethanolamine-N-[poly(ethylene glycol)2000-N'-carboxyfluorescein] (DSPE-PEG2k-FITC, Avanti catalog no: 810120C), and 0.3% ATTO 647-labeled 1,2-dioleoyl-sn-glycero-3-phosphoethanolamine (ATTO 647-DOPE, ATTO-TEC catalog no: AD 647-161). Lipids were spread on an indium tin oxide (ITO)-coated microscope slide (Diamond Coatings, 8-12 Ohm slide) and dried under vacuum for 30 minutes.

A 2 mm rubber gasket was sandwiched between the ITO-coated slide containing lipids and a clean ITO-coated slide, and the interstitial space filled with 75 mM sucrose solution. A sinusoidal electric potential of amplitude 3V (peak-to-peak) and frequency 10 Hz was applied to the chamber for two hours at 50 °C. After two hours frequency was changed to 2 Hz for 30 minutes. The resulting GUVs were collected, stored at room temperature and used within one day.

#### D. Surface preparation

Glass cover slips No. 1.5 (Fisher) were cleaned with piranha solution (3:1 sulfuric acid:hydrogen peroxide) for five minutes and then washed with deionized water. The cover slips were then made hydrophobic via reaction with trimethylchlorosilane (Sigma) vapors in a vacuum chamber, under house vacuum for ten minutes. A 6 mm cylindrical polydimethylsiloxane (PDMS) chamber was attached to the cover slip surface to hold liquids. Cover slips were incubated with 200 nM heavy meromyosin (HMM) for five minutes. After five minutes, 0.1 mg/mL polylysine-grafted-PEG (PLL-g-PEG) was added and the HMM/PLL-g-PEG solution incubated for another five minutes. The coverslip was then washed, first with A-buffer, and then with MilliQ water.

#### E. Assembling actomyosin cortex on a lipid bilayer

GUVs in MilliQ water were added to the cover slip chamber. The cover slip was heated to 37°C for at least 20 minutes, during which time GUVs ruptured on the treated surface. Unbound GUVs were then washed from the cover slip with A-buffer. For experiments with activity, the cover slip was then incubated in 1 µM F-actin for 10 minutes at 37 °C. F-actin spontaneously adsorbed to the liquid-ordered phase of lipid bilayer, presumably via electrostatic attraction to DOTAP. Unbound actin was washed from the cover slip with A-buffer. The cover slip with lipids and actin was incubated in 500 nM Myosin II for ten minutes at 37°C before the unbound myosin II washed away with A-buffer.

#### F. Microscope for all imaging experiments

All imaging was carried out on an inverted Nikon Ti2-Eclipse microscope (Nikon Instruments) using an oil-immersion objective (Apo 100x, NA 1.45, oil). Lumencor SpectraX Multi-Line LED Light Source was used for excitation

(Lumencor, Inc). Fluorescent light was spectrally filtered with emission filters (432/36, 515/30, 595/31, and 680/42; Semrock, IDEX Health and Science) and imaged on a Photometrics Prime 95 CMOS Camera (Teledyne Photometrics). Microscope images were collected using MicroManager 1.4 software [5].

#### G. Imaging passive domain growth

Planar lipid membranes without actin were gently heated with a hair dryer until the domains melted, as confirmed by fluorescence microscopy. The hair dryer was then withdrawn, and the bilayer imaged over time as it cooled and the domains re-formed and grew.

#### H. Imaging active domain growth

Due to the temperature-sensitivity of the actin-membrane interactions, actomyosin cortex, and DMNPE-caged ATP, active domain growth was tracked at room temperature. Thus, domains were tracked as they grew from a small initial size, fixed by the actin network, to a larger size under the influence of actomyosin activity.

Photosensitive caged ATP (1µM DMNPE-caged ATP, Invitrogen catalog no: A1049) was added to planar lipid membranes with actin and myosin before imaging. The sample was irradiated with a brief (<1 s) pulse of 404 nm light, converting the caged ATP to usable ATP. Domains and actin were then imaged over time, as the actomyosin contracted, and the domain size and morphology evolved.

#### I. Image analysis

Microscope images of lipid domains were thresholded in ImageJ and the area and perimeter of the resulting black and white images calculated using MATLAB (MathWorks). Characteristic domain size $a$ was taken to be the ratio of area/perimeter. Domain size versus time was fitted to an equation of the form:

$$a = (At + B)^C \tag{1}$$

where the prefactor $A$ is related to solvent viscosity, substrate friction, and domain area fraction (Eq. 21); $B$ is related to the initial size at $t = 0$, $a_0 = B^C$; and $C$ is the growth rate exponent. For plots of domain size versus time in Fig. 2 of the main text, time is shifted for each experimental realization, so that $t = 0$ corresponds to the point at which $a = 0$. Thus, $t_{\text{plot}} = t + \frac{B}{A}$ where $t = 0$ is the time at which imaging begins, and $t_{\text{plot}} = 0$ is the time at which domains begin to form.

### III. THEORY OF DOMAIN GROWTH

In this section we discuss multiple theoretical approaches to predict the growth rate of lipid domains. We first derive the domain growth rate via a Cahn-Hilliard field-based approach in which we describe the evolution of an order parameter $\phi$, as described by Lifshitz and Slyozov [6, 7]. This approach assumes that domains grow solely via molecular transport, i.e. an evaporation/condensation of Ostwald ripening mechanism.

We then derive the rate of domain growth via a Smoluchowski agent-based approach, in which we treat the domains as individual particles [8]. By treating particle coalescence as a severely transport-limited two-species reaction, we show that domains grow at the same rate via coalescence, as they do via evaporation/condensation. Finally, by applying a simple shear flow to this Smoluchowski-based description, we demonstrate that 2D flows can increase the rate of domain coarsening.

#### A. Lifshitz-Slyozov Scaling

Consider a droplet of radius $R$ and concentration, $\phi = +1$, surrounded by bulk concentration phase, $\phi = -1$. The interface has surface tension $\sigma$. In the bulk phase, there is a fluctuation $\varepsilon(r)$ because the concentration is not uniformly $\phi = -1$ near $r = R$. Assume $\phi(r) = -1 + \varepsilon(r)$, $\varepsilon \ll 1$, and the gradients of order parameter in the bulk phase is negligible. The Cahn-Hilliard model now becomes $\partial_t \varepsilon = 2\nabla^2\varepsilon + \mathcal{O}(\varepsilon^2)$. Assuming that the fluctuations quickly reach a steady state at all times compared to the growth of domains (quasi-static approximation):

$$\nabla^2 \varepsilon = 0 \ . \tag{2}$$

Thus, the fluctuation field is defined by

$$\varepsilon(r) = C_1 \ln(r) + C_2 \tag{3}$$

where $C_1$ and $C_2$ are integration constants. Applying the boundary conditions $\varepsilon(r = R) = \sigma/(2R)$, $\varepsilon(r = L) = 0$ where $L$ is the 2D box length, and $L \gg R$ gives

$$\therefore \ \varepsilon(r) = \frac{\sigma}{2R} \frac{\ln(r/L)}{\ln(R/L)} \ . \tag{4}$$

The droplet radius grows according to $\mathrm{d}R/\mathrm{d}t$. Performing a flux balance at the interface gives

$$j = -2\nabla\varepsilon|_{r=R} = \Delta\phi \frac{\mathrm{d}R}{\mathrm{d}t} \tag{5}$$

where $j$ is the surface flux and $\Delta\phi$ is the difference in order parameter across the interface. Thus,

$$\frac{\mathrm{d}R}{\mathrm{d}t} = \frac{\sigma}{2R^2 \ln(R/L)} \ . \tag{6}$$

Solving this first order differential equation yields

$$R^3 \left[ \ln\left(\frac{R}{L}\right) - 1 \right] = \frac{3}{2}\sigma t + C \tag{7}$$

where $C$ is an integration constant. The area fraction of domains in the 2D box is $\pi(R/L)^2$, so in the case of constant domain area fraction, $R/L$ is constant. If the initial droplet radius is zero, then $R \sim t^{1/3}$.

#### B. Smoluchowski Coalescence

Domain growth via coalescence can be described using a Smoluchowski coagulation model, which is typically used to model the flocculation of colloids of a fixed size. Here we first adapt the Smoluchowski description of colloid flocculation to calculate the growth rate of singlet domains in the absence of flow (passive). This approach has been previously applied to lipid domain growth by Frolov et al. [9], and treats domains as particles of uniform size, which merge infinitely quickly and irreversibly upon contact. We then apply a simple shear flow to the 2D system and find the dependence of the domain growth rate on the colloidal Péclet number $\mathrm{Pe_c}$, as described by Russel, Saville, and Schowalter [8] for 3D colloidal aggregation.

##### 1. *Passive domain growth in the absence of flow*

To model the growth of 2D passive domains in the absence of flow, we consider two identical circular particles (particles 1 and 2) of size $a$ and colloidal scale diffusivity $D_\mathrm{c}$. These particles represent singlet lipid domains. This pair of particles has the pair distribution function $g(\mathbf{r}, t)$, which evolves according to the equation

$$\frac{\partial g}{\partial t} + \nabla_x \cdot \mathbf{j}_1 + \nabla \cdot (\mathbf{j}_1 - \mathbf{j}_2) = 0 \tag{8}$$

where $\mathbf{j}_i$ denotes the flux of species $i$, $\nabla_x$ is the gradient operator relative to the origin, and $\nabla_r$ is the gradient operator relative to the center of particle 1. Assuming spherical particles and spatial invariance reduces the Smoluchowski equation to

$$\frac{\partial g}{\partial t} + \nabla_r \cdot \mathbf{j}_\mathrm{rel} = 0 \tag{9}$$

where $\mathbf{j}_{\rm rel} = \mathbf{j}_1 - \mathbf{j}_2$. We define $\mathbf{j}_{\rm rel}$ as

$$\mathbf{j}_{\rm rel} = \mathbf{U}g - D_{\rm c}\left(\nabla_r g + g\frac{\nabla_r V}{k_{\rm B}T}\right) \tag{10}$$

where $\mathbf{U}$ is bulk fluid velocity and $V$ is the inter-particle potential.

$$\mathbf{j}_{\rm rel} = -D_{\rm c}\left(\nabla_r g + g\frac{\nabla_r V}{k_{\rm B}T}\right) \tag{11}$$

for all $r \geq 2a$.

Integrating Eq. 9 over all space gives

$$\frac{\partial n}{\partial t} + \int \nabla_r \cdot \mathbf{j}_{\rm rel}\ {\rm d}\mathbf{r} = 0 \tag{12}$$

where $n$ is the number density of singlet domains, defined as $n(t) = \int g(\mathbf{r}, t)\ {\rm d}\mathbf{r}$. Applying the divergence theorem converts the integral over all space in Eq. 12 into a line integral over the perimeter of domain 1:

$$\int \nabla_r \cdot \mathbf{j}_{\rm rel}\ {\rm d}\mathbf{r} = \oint_{r=2a} \mathbf{n}\cdot\mathbf{j}_{\rm rel}\ {\rm d}S \tag{13}$$

where $\mathbf{n}$ is the normal vector. Assuming a steady state flux of domains at the surface of species 1 makes Eq. 13 equal to a constant $J = \oint_{r=2a} \mathbf{n}\cdot\mathbf{j}_{\rm rel}\ {\rm d}S$. Thus the number density of domains evolves in time according to

$$\frac{\partial n}{\partial t} = -J\ . \tag{14}$$

To find $J$, we solve Eq. 9, which reduces to

$$\frac{\rm d}{{\rm d}r}\left[r\left(\frac{{\rm d}g}{{\rm d}r} + g\frac{{\rm d}V}{{\rm d}r}\right)\right] = 0 \tag{15}$$

for a radially symmetric system at steady state, with a flux defined by Eq. 11. Applying the boundary conditions $g(r = 2a) = 0$ and $g(r = L) = n^2$ for a large, but finite system size $L$ gives the solution

$$g(r) = \frac{n^2 e^{-V/k_{\rm B}T}\int_{2a}^{r}\frac{e^{-V/k_{\rm B}T}}{r}{\rm d}r}{\int_{2a}^{L}\frac{e^{-V/k_{\rm B}T}}{r}{\rm d}r} \tag{16}$$

Thus,

$$J = 4\pi a D_{\rm c}\left.\frac{{\rm d}g}{{\rm d}r}\right|_{r=2a} = \frac{2\pi D_{\rm c} n^2}{\ln\left(\frac{L}{2a}\right)} \tag{17}$$

Assuming a constant domain area fraction $\phi_{\rm A}$ of $N$ circular domains,

$$J = \frac{2\pi D_{\rm c} n^2}{\ln\left(\frac{1}{2}\sqrt{\frac{N\pi}{\phi_{\rm A}}}\right)}\ . \tag{18}$$

Substituting Eq. 18 into Eq. 14 and applying the diffusivity $D_{\rm c} = \frac{k_{\rm B}T}{16\eta a}$ for 2D domains much larger than the Saffman-Delbrück limit, $a \gg L_{\rm SD}$ [10], gives

$$\frac{{\rm d}n}{{\rm d}t} = -\frac{k}{a}n^2 \tag{19}$$

where $k = \frac{\pi k_{\rm B}T}{8\eta\ln\left(\frac{1}{2}\sqrt{\frac{N\pi}{\phi_{\rm A}}}\right)}$ and $\eta$ is the 3D solvent viscosity.

At constant area fraction $\phi_{\rm A} \sim na^2$, this differential equation can be rewritten in terms of domain size $a$

$$\frac{{\rm d}a}{{\rm d}t} = \frac{2k\phi_{\rm A}}{a^2}\ . \tag{20}$$

Therefore, domains grow according to the relation

$$a = \left(6k\phi_{\rm A}t + a_0^3\right)^{1/3} \tag{21}$$

where $a_0$ is the initial domain size. Thus, domains that grow from $a_0 = 0$, purely by coalescence, follow the scaling

$$a \sim t^{1/3} \tag{22}$$

*2. Domain growth in simple shear flow*

Starting from Eq. 9, the flux now contains a velocity term, $\mathbf{U} = \dot{\gamma} r \mathbf{e_r}$, where $\dot{\gamma}$ is the shear rate.

$$\mathbf{j}_{\text{rel}} = \mathbf{U}g - D_{\text{c}}\left(\nabla_r g + g\frac{\nabla_r V}{k_{\text{B}}T}\right) \tag{23}$$

The boundary conditions are the same as before. In non-dimensional form, the Smoluchowski equation is

$$\frac{\partial g}{\partial t} + \text{Pe}_{\text{c}}\nabla_r \cdot \mathbf{r}g - \nabla_r \cdot \left(\nabla_r g + g\frac{\nabla_r V}{k_{\text{B}}T}\right) = 0 \tag{24}$$

The colloidal Péclet number is $\text{Pe}_{\text{c}} = \dot{\gamma}a^2/D$, and is distinct from the Péclet number Pe used in the simulations, as its characteristic length scale $a$ and diffusivity $D$ correspond to whole domains, and are not molecular (lipid) parameters. To solve the equation, it is assumed that $g$ can be expanded in powers of $\text{Pe}_{\text{c}}$ as $g = n^2(g_0 + \text{Pe}_{\text{c}}\, g_1 + \mathcal{O}({\text{Pe}_{\text{c}}}^2))$ in the limit $\text{Pe}_{\text{c}} \ll 1$. The leading order term, $g_0$, is the solution for the passive case.

$$g_0 = \frac{\ln(r/2a)}{\ln(L/2a)} \tag{25}$$

The steady-state behavior far away reveals that there is a boundary layer.

$$\underbrace{\nabla_r \cdot \left(\nabla_r g_0 + g_0\frac{\nabla_r V}{k_{\text{B}}T}\right)}_{\frac{1}{r}\sim 0} + \text{Pe}_{\text{c}}\nabla_r \cdot \left(\nabla_r g_1 + g_0\frac{\nabla_r V}{k_{\text{B}}T}\right) = \text{Pe}_{\text{c}}\nabla_r \cdot \underbrace{\mathbf{r}g_0}_{r\frac{\ln(r/2a)}{\ln(L/2a)}\sim\infty} + \mathcal{O}({\text{Pe}_{\text{c}}}^2) \tag{26}$$

As $r \to \infty$, convection dominates diffusion in the small $\text{Pe}_{\text{c}}$ limit. By matching the order of both terms, we get $\text{Pe}_{\text{c}}\, r^2 \ln(r/2a)/\ln(L/2a) \sim 1$. Following a similar argument as before, $L/2a$ is the inverse area fraction, which is constant due to mass conservation. We argue that for large $r$ and very small $\text{Pe}_{\text{c}}$ such that $L \gg 1/\sqrt{\text{Pe}_{\text{c}}}$, we can approximate $\text{Pe}_{\text{c}}\, r^2 \ln r \approx \text{Pe}_{\text{c}}\, r^2 \sim 1$. Thus, the boundary layer region is re-scaled as $r \sim {\text{Pe}_{\text{c}}}^{-1/2}$.

After performing a singular perturbation analysis at the boundary layer, and asymptotically matching with the solution at small $r$, we get an expression for flux.

$$J = \frac{2\pi k_{\text{B}}T}{16\eta a \ln\left(\frac{1}{2}\sqrt{\frac{N\pi}{\phi_{\text{A}}}}\right)} n^2\left(1 + C{\text{Pe}_{\text{c}}}^{1/2}\right) \tag{27}$$

The constant $C$ is a numerical factor obtained after asymptotic matching. Substituting this in Eq. 14, and using $n = \phi_{\text{A}}/(\pi a^2)$ gives

$$a^2\frac{\text{d}a}{\text{d}t} = k(1 + C{\text{Pe}_{\text{c}}}^{1/2}) \tag{28}$$

The Péclet number depends on $a$ as $\text{Pe}_{\text{c}} = \dot{\gamma}a^2/D_{\text{c}} = 16\dot{\gamma}\eta a^3/(k_{\text{B}}T)$. The solution to the differential equation is

$$\frac{2}{3C_1^2}[C_1 a^{3/2} - \ln(1 + C_1 a^{3/2})] = C_0 + kt \tag{29}$$

Here, $C_1 = 16C\dot{\gamma}\eta/(k_{\text{B}}T)$, and $C_0 = (2/3C_1^2)(C_1 a_0^{3/2} - \ln(1 + C_1 a_0^{3/2}))$. For small $a$, we can expand $\ln(1 + C_1 a^{3/2})$ as a power series.

$$\frac{2}{3}\left[\frac{a^3}{2} + \mathcal{O}(a^{9/2})\right] = C_0 + kt \implies a \sim t^{1/3} \tag{30}$$

We see that passive growth dominates during early time growth. Similarly, for large $a$, we get

$$\frac{2}{3}\left[\frac{a^{3/2}}{C_1} - \frac{\ln(C_1 a^{3/2})}{C_1} + \mathcal{O}(a^{-3/2})\right] = C_0 + kt \implies a \sim t^{2/3} \tag{31}$$

The $a^{3/2}$ term dominates the log terms in the expansion, so effectively, the domain size scales as $t^{2/3}$ at long times.

### IV. SIMULATION DETAILS

The Cahn-Hilliard model is solved numerically using pseudo-spectral methods. The evolution of the Fourier transformed order parameter $\phi_k$ is

$$\frac{\mathrm{d}\phi_k}{\mathrm{d}t} + \mathcal{F}[\mathbf{v}\cdot\nabla\phi] = -Mk^2\mathcal{F}\left[\frac{\delta f}{\delta\phi}\right] - M\kappa k^4\phi_k. \tag{32}$$

This first-order differential equation is solved using forward finite difference. The higher order gradient terms ($\nabla^4$) are treated implicitly. The bulk free energy, $\delta f/\delta\phi = \phi^3 - \phi$, is a non-linear term so a 2/3 anti-alias filter is applied to remove high-frequency noise.

The initial system configuration is $\phi = \phi_{\mathrm{avg}} + 0.1 \times \text{white noise}$. For the main text figures, the average concentration is $\phi_{\mathrm{avg}} = 0.3$. The simulation box is a $512^2$ grid with a box length of $32\pi$. The time step is controlled with adaptive time stepping, $\Delta t = [\mathrm{tol}/(\phi(t;\Delta t) - \phi(t;\Delta t/2)]^{1/5}$, where the tolerance, tol is $10^{-4}$.

#### A. Metric for domain size

In the main text, the domain size is calculated using area/perimeter. There is an alternate metric based on the structure factor of domains, $S(k,t) = \langle\phi_k(t)\phi_{-k}(t)\rangle$, that also yields similar results. In this metric, the domain size is the inverse of the first moment of the structure factor:

$$a(t) = \frac{\int_{k_{\mathrm{min}}}^{k_{\mathrm{cut}}} S(k,t)\mathrm{d}k}{\int_{k_{\mathrm{min}}}^{k_{\mathrm{cut}}} kS(k,t)\mathrm{d}k}. \tag{33}$$

Here $k_{\mathrm{min}}$ is the minimum non-zero wave-number,$k_{\mathrm{cut}} = 2k_{\mathrm{max}}$ is the cut-off wavelength, and $k_{\mathrm{max}}$ is the wave number corresponding to the maximum of the structure factor. Fig. S1 shows that using the area/perimeter ratio of domains yields similar scaling for $a(t)$ as the first moment of the static structure factor.

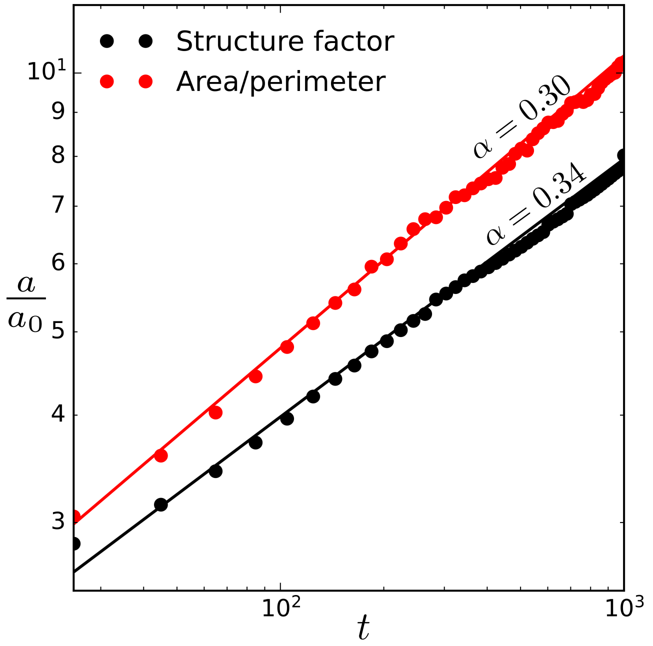


FIG. S1. Comparison of area/perimeter and structure factor as metrics for domain size for passive coarsening. Domain growth scales similarly regardless of the metric used to quantify domain size. The simulation parameters are ($M$, $\kappa$)=(1, 0.25).

[1] J. A. Spudich and S. Watt, The Regulation of Rabbit Skeletal Muscle Contraction, Journal of Biological Chemistry **246**, 4866 (1971).

[2] S. MacLean-Fletcher and T. D. Pollard, Identification of a factor in conventional muscle actin preparations which inhibits actin filament self-association, Biochemical and Biophysical Research Communications **96**, 18 (1980).

[3] S. S. Margossian and S. Lowey, Preparation of myosin and its subfragments from rabbit skeletal muscle, in *Methods in Enzymology*, Vol. 85 (1982) pp. 55–71.

[4] M. I. Angelova and D. S. Dimitrov, Liposome electroformation, Faraday Discussions of the Chemical Society **81**, 303 (1986).

[5] A. D. Edelstein, M. A. Tsuchida, N. Amodaj, H. Pinkard, R. D. Vale, and N. Stuurman, Advanced methods of microscope control using $\mu$Manager software, Journal of Biological Methods **1**, e10 (2014).

[6] I. Lifshitz and V. Slyozov, The kinetics of precipitation from supersaturated solid solutions, Journal of Physics and Chemistry of Solids **19**, 35 (1961).

[7] A. J. Bray and C. L. Emmott, Lifshitz-Slyozov Scaling For Late-Stage Coarsening With An Order-Parameter-Dependent Mobility, Phys. Rev. B **52**, R685 (1995).

[8] W. B. Russel, D. A. Saville, and W. R. Schowalter, *Colloidal Dispersions* (Cambridge University Press, 1989).

[9] V. Frolov, Y. Chizmadzhev, F. Cohen, and J. Zimmerberg, "Entropic Traps" in the Kinetics of Phase Separation in Multicomponent Membranes Stabilize Nanodomains, Biophysical Journal **91**, 189 (2006).

[10] C. A. Stanich, A. R. Honerkamp-Smith, G. G. Putzel, C. S. Warth, A. K. Lamprecht, P. Mandal, E. Mann, T. A. D. Hua, and S. L. Keller, Coarsening dynamics of domains in lipid membranes, Biophysical Journal **105**, 444 (2013).
